# TNFR2 Agonism as a Sex-Specific Therapy for Novel Osteoarthritis-Induced Cardiac Dysfunction

**DOI:** 10.64898/2026.07.06.736778

**Authors:** Pranav Prasoon, Kelly Tammen, Rebekah Russo, Aravind Meyyappan, Manushri Dalvi, Roman Fischer, Melanie Eschborn, Sreejita Arnab, Luke Brabbee, Lauren Schneider, Kayla Nguyen, David Mendelowitz, Matthew W Kay, John R. Bethea

## Abstract

Osteoarthritis (OA), a degenerative joint disease, is associated with increased systemic inflammation, chronic pain, and cardiovascular dysfunction. Epidemiological evidence establishes that OA increases the risk of cardiovascular disease (CVD) threefold, yet the causal role of OA’s contributions remains underexamined. We assessed cardiac function longitudinally following destabilization of the medial meniscus (DMM) surgery to induce osteoarthritis in mice. DMM-mice exhibited significant, sexually dimorphic alterations in echocardiographic parameters. Female DMM mice developed impaired relaxation with altered E/A ratios, increased E/e’ ratios, and prolonged intraventricular relaxation time with no change in ejection fraction, while male DMM mice showed progressive systolic dysfunction with decreasing ejection fraction, increased E/e’ ratio, and prolonged intraventricular contraction time. Transcriptomic profiles and biochemical analyses demonstrated divergent cellular responses involving fibrosis and oxidative stress in female mice, whereas autophagic and apoptotic responses were observed in male mice. Using a tumor necrosis factor 2 (TNFR2) agonist shown to reduce systemic inflammation, we investigated its potential therapeutic role in the context of OA-induced cardiovascular dysfunction. TNFR2 agonism proved to be effective both prophylactically and therapeutically for female diastolic dysfunction. While prophylactic and therapeutic administration delayed male systolic dysfunction, the efficacy declined over time. Our findings demonstrate evidence of a novel sexually dimorphic model of OA-induced CVD that recapitulates the sexually dimorphic pattern of patient phenotypes and a promising new therapeutic approach to CVD.

**Translational Relevance:** Osteoarthritis patients have higher, often unrecognized, cardiovascular risk, yet preclinical models linking joint disease to cardiac dysfunction remain unexplored. Using a murine preclinical model of OA reveals the key findings. First, OA alone drives sex-specific cardiac phenotypes - females develop diastolic dysfunction, whereas males develop progressive systolic impairment. Second, selective TNFR2 agonism prevents and reverses OA-induced diastolic dysfunction in female mice and delays systolic decline in males. These findings suggest sex-dependent cardiac monitoring in OA patients and indicate that TNFR2-targeted therapy will likely be a sex-informed intervention to provide cardioprotective benefit.

**Graphical Abstract:** 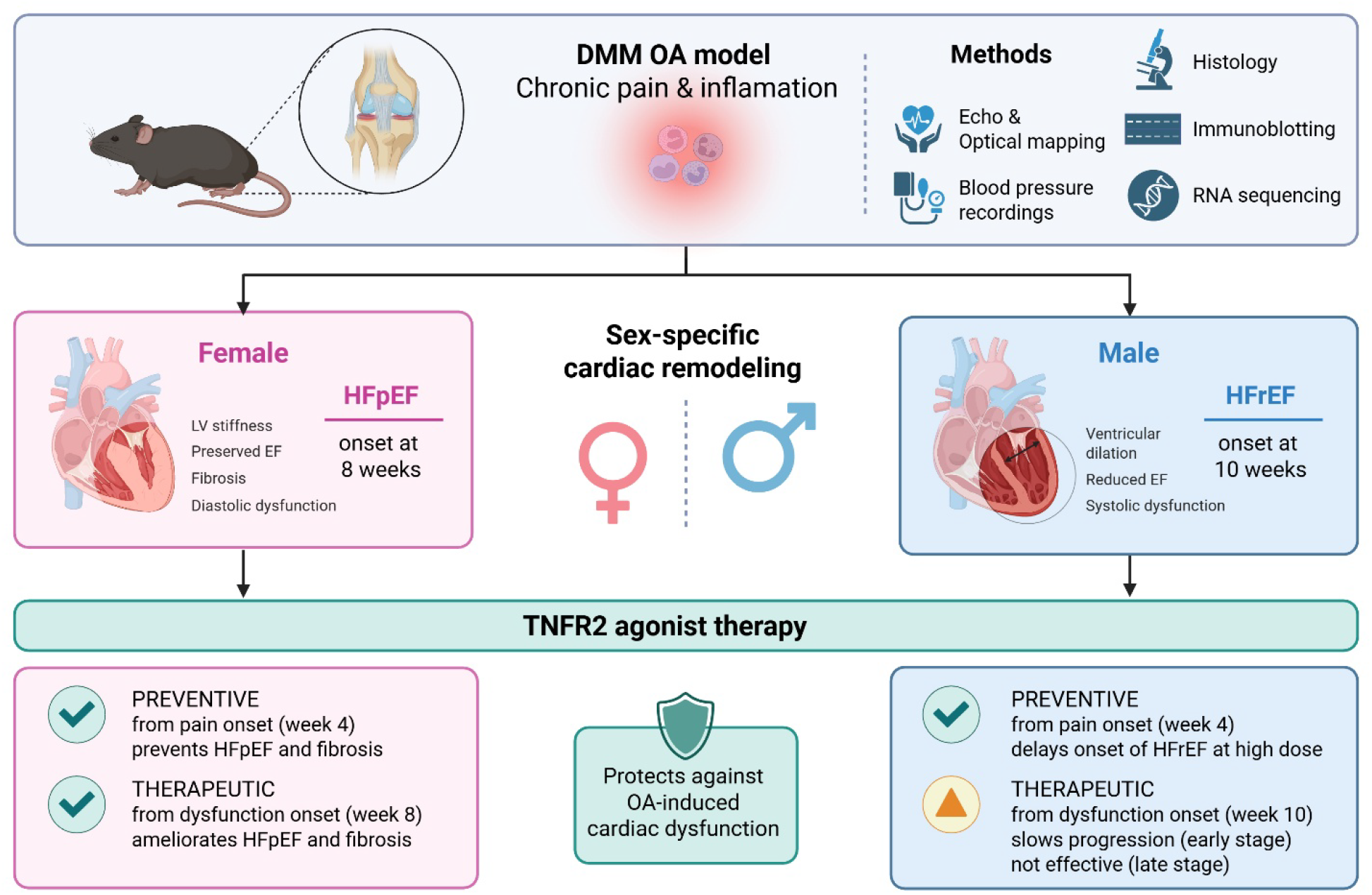

## Introduction

Mounting evidence indicates chronic inflammation contributes to the development and progression of heart failure (HF) pathophysiology^1–5^. HF, a leading cause of morbidity and mortality globally, affects over 56.2 million individuals with a 5-year survival rate below 50% following diagnosis. Osteoarthritis (OA), a chronic inflammatory joint disease, affects over 500 million people worldwide and is among several conditions with increased HF incidence^7,8^. Substantial epidemiological evidence establishes that OA patients have a threefold increased risk of cardiovascular disease (CVD), predominantly HF, compared to healthy age-matched individuals^9–15^. Heart failure with preserved ejection fraction (HFpEF) (LVEF ≥50%) accounts for approximately 50% of HF cases and disproportionately affects postmenopausal women, whereas heart failure with reduced ejection fraction (HFrEF) (LVEF ≤40%) predominantly affects men^16–19^. Previous studies show that an elevated incidence of HF is associated with persistent low-grade systemic inflammation, leading to metabolic and endothelial dysfunction in HFpEF and cardiomyocyte loss in HFrEF, accelerating pathological cardiac remodeling ^12,15,20,21^. Despite the strong epidemiological association, the causality and contribution of OA-driven inflammation to HF progression remain poorly defined, partly due to a lack of appropriate preclinical models. To address this gap, we examined the development and progression of cardiac dysfunction in an established OA murine model: surgical destabilization of the medial meniscus (DMM).

Despite established HFrEF therapies, including angiotensin receptor-neprilysin inhibitors (ARNI), beta-blockers, mineralocorticoid receptor antagonists (MRA), and sodium-glucose cotransporter 2 inhibitors (SGLT2i), HFpEF treatment options remain limited. Empagliflozin, currently the only FDA-approved HFpEF therapy, reduces hospitalizations, but provides only modest mortality benefits, reflecting the heterogeneous and multifactorial nature of HFpEF pathophysiology ^22,23^. Clinical trials targeting inflammatory cytokines (TNF, IL-1, IL-6) in HF show limited efficacy ^2,24–28^. We postulate that this failure reflects the heterogeneous nature of cardiovascular inflammation and non-selective inhibition of both protective and pathological signaling. TNFR1 activation promotes proinflammatory and pathological responses, while TNFR2 signaling is cardioprotective ^24,29–31^. Current anti-TNF therapies block TNFɑ binding to both TNFR1 and TNFR2, potentially negating beneficial TNFR2 signaling^25^.

Selective TNFR2 activation represents a promising therapeutic strategy. Using cell-specific genetic and pharmacological approaches, we and others showed that TNFR2 activation mitigates pathological inflammation across preclinical models in both sexes ^31–38, 40, 63^. Here, we identified a novel non-genetic model of OA-induced cardiac dysfunction and investigated the therapeutic potential of a selective TNFR2 agonist.

## Materials and Methods

### Animals

Eleven-week-old male and female C57BL/6J (Cat. 000664) mice were obtained from The Jackson Laboratory. Before the experiments, mice were given a one-week acclimation period. Animals were housed under a 12-hour light/dark cycle with access to food and water *ad libitum*. Upon study termination at 16 weeks (females) and 24 weeks (males), mice were anesthetized using a ketamine and xylazine cocktail prior to perfusion. The Institutional Animal Care and Use Committee (IACUC) at George Washington University (GWU), Washington, DC, USA approved these animal experiments. They were conducted in accordance with the animal use protocol (GWU IACUC no. A2024-091) and the guidelines of the National Institutes of Health. All ethical guidelines were adhered to during the execution of this study.

### Osteoarthritis (OA)-induced Heart Failure model

Destabilization of the medial meniscus (DMM) surgically induced OA. 12-week-old C57BL/6J male and female mice underwent DMM as previously described ^39^. Mice were anesthetized with Isoflurane (1–1.5% in oxygen) administered via mask induction throughout the surgical procedure, and the right knee joint was shaved and sterilized with Betadine and ethanol. A medial parapatellar incision exposed the joint capsule, and the medial meniscotibial ligament (MMTL) was transected under a surgical microscope to destabilize the medial meniscus. The capsule and skin were sutured using 6-0 absorbable sutures and surgical glue, respectively. Buprenorphine, 0.05 mg/kg, was administered subcutaneously as needed for up to 48 hours post-operatively. Mice were monitored daily for signs of distress and allowed unrestricted activity until sacrifice. At the end of the experiment, mice were euthanized via intraperitoneal injection of ketamine/xylazine (100:10 mg/kg, i.p, single injection) in accordance with institutional guidelines, and the tissue was extracted for further analysis.

### Drug

A selective murine TNFR2 agonist (TNFR2 Ag) was employed in this study, composed of a covalently stabilized, TNFR2-specific single-chain TNF trimer fused to an effector-deficient Fc domain^40^. Four weeks following DMM surgery, a preventive TNFR2 Ag was administered intraperitoneally twice weekly in both male (1 mg/kg and 10 mg/kg) and female (1 mg/kg) mice to evaluate its preventive effects on behavioral and cardiac function tests. We assessed the therapeutic effects of TNFR2 Ag (I.P., administered twice weekly) on cardiac function following DMM surgery at 10 weeks (male) and 8 weeks (female) at 10 mg/kg and 1 mg/kg, respectively.

### Von Frey Testing

The Von Frey test assessed mechanical allodynia as previously described ^41,42^. Mice were individually placed in plexiglass chambers elevated on a mesh-wire platform and permitted to acclimate for 45 to 60 minutes prior to testing. Using the up-down method ^43^, von Frey filaments (Touch-test sensory evaluator) of varying diameters (force range: .02g-2g) stimulated the plantar surface of the hind paw ipsilateral to the surgical site. In each trial, an intermediate filament weighing 0.16 g was applied perpendicular to the skin, causing a slight bend for 3 seconds. A positive response hindpaw withdrawal response was only marked as a pain response if paired with at least one of the following behaviors: stimulus avoidance behavior, tucking the tail under the stimulated hind paw, and/or licking the hind paw, indicating cognitive recognition of the stimulus as being painful. If no response occurred, the next larger filament was tested, and a series of 5 trials was completed once the first positive pain response was noted. The statistical method outlined by Dixon (1965) was used to calculate the 50% mechanical withdrawal threshold. The Baseline withdrawal threshold of the mice was assessed one day prior to DMM surgery, and weekly von Frey testing was conducted from 3 to 12 weeks post-DMM surgery.

### Echocardiography (ECHO)

ECHO measurements were acquired using a VisualSonics Vevo 3100 while recording body temperature, respiratory rate, and the ECG. LV outflow tract diameter was measured using a 2D parasternal long-axis view. The LV diastolic posterior wall diameter (LVPWd) and interventricular septal diameter (IVSd), LV diameter in diastole (LVDd) and systole (LVDs), and LV systolic and diastolic posterior-wall (LVPWs,d) thickness were measured using an M-mode parasternal short-axis view at the plane of the papillary muscles. These measurements were used to calculate fractional shortening (FS) and LV ejection fraction (EF%). Diastolic function was assessed using blood flow velocity through the mitral valves, including early diastolic mitral inflow (E-wave), late diastolic mitral inflow (A-wave), and early-to-late diastolic mitral inflow ratio (E/A ratio), which were measured using an apical four-chamber view. The longitudinal velocity of the myocardium (e’) was assessed using tissue doppler imaging at the mitral annulus. Longitudinal strain, radial strain, circumferential strain, and outer wall delay were assessed using a strain analysis of B-mode images from the parasternal long-axis view and short axis views. HR and cardiac output were measured using the blood velocity and pressure in the ascending aorta provided by continuous-wave Doppler imaging^44^.

### Blood pressure recordings

Blood pressure measurements-systolic blood pressure (SBP), diastolic blood pressure (DBP), and mean arterial pressure (MAP)-were recorded with CODA high throughput tail cuff system (CODA High Throughput System, Noninvasive Blood Pressure System, CODA-HT8; Kent Scientific Corp., Torrington, Connecticut). Conscious mice were a placed on a heated platform (33-34°C), covered with a blanket, and in restraint tubes for duration of acclimation and recording. Restrained mice rested quietly in the room for 5 minutes prior to recording. The same person performed blood pressure measurements between 7-11 am. Mice were acclimated to the system for two days prior to recording. Values from 10-20 cycles were used to calculate the mean SBP, DBP, MAP, and standard deviation for each mouse across accepted recordings. 7 acclimation cycles were used for each test with 20 recordings following. Animals were excluded from analysis if they had fewer than 10 accepted recordings ^45^.

### Histology

For tissue used in histology, mice were perfused using the aorta with PBS then 10% Neutral Buffered Formalin (NBF) and kept on ice and drop-fixed in 10% NBF for 72 hours. All stained sections were examined using a Leica DMR microscope with a 10x objective, equipped with a Leica DFC 300 FX digital camera.

#### Knee

Samples were decalcified with 14% EDTA for 2 weeks. Tissues were then transferred to 30% sucrose solution for 3 days then embedded in optimal cutting temperature (O.C.T.) compound. O.C.T embedded tissues were sectioned at 10 μm sections and stained with safranin O-fast green. Proteoglycan loss and fibrillation of knee tissue in knee joints indicated cartilage degradation. Histological scoring was conducted on four to six sections per animal.

#### Heart

Tissues were transferred to 30% sucrose solution for 3 days then embedded in O.C.T. for sectioning. O.C.T embedded tissues were sectioned at 15 μm sections and stained with Hematoxylin and Eosin (H&E). Protocol was obtained and adapted to heart tissue. Histological scoring was conducted on 3 sections per animal.

### Histological Scoring (Knee and Heart)

#### Knee

We utilized the OARSI scoring method, a semiquantitative scoring system for murine histopathology, for the knee sections. All four quadrants of the knee joint were evaluated: medial femoral condyle (MFC), lateral femoral condyle (LFC), medial tibial plateau (MTP), and lateral tibial plateau (LTP). A score from 0 to 6 was given to each quadrant of 3 serial sections per animal, 12 total values per animal. The final histological scores were the average of all the individual values, to calculate the average summed score for each experimental group. Two blinded observers scored all the sections. A third observer was involved if scores differed by greater than one point on the OARSI scale ^46^.

#### Heart

Left ventricular wall thickness was measured using ImageJ software. Three measurements of the left ventricular wall thickness and distance between the intraventricular septum and left ventricular wall were taken per sample and averaged with three animals per group.

### RNA sequencing and Gene Ontology analysis

Left ventricular RNA was isolated from male and female mice from all treatment groups: naïve, vehicle-treated OA, or TNFR2 agonist-treated OA groups. Stranded mRNA libraries were prepared using the Illumina TruSeq stranded mRNA kit and sequenced on an Illumina NovaSeq platform. Reads were quality-filtered, adapter-trimmed, and depleted of ribosomal and mitochondrial contaminants before alignment to the mouse genome (GRCm38, Ensembl annotation) using STAR v2.7.3a. Gene-level counts were quantified with HTSeq v0.11.2 and analyzed in Rstudio using DESeq2. Counts were normalized using DESeq2 library, and differential expression was tested using the Wald test with Benjamini–Hochberg correction. Genes that met p-value < 0.05 and absolute fold change ≥ 2 were carried forward as differentially expressed genes.

Functional enrichment analysis was performed using clusterProfiler and org.Mm.eg.db. Differentially expressed gene symbols were converted to Entrez IDs with AnnotationDbi and tested for enrichment across Gene Ontology Biological Process, Cellular Component, and Molecular Function categories. Enrichment results were visualized as dot plots, and gene–term relationships among enriched pathways were displayed using cnet plots.

### Western Blots

For fresh tissue used in Western blots, animals were perfused through the aorta with PBS to remove blood. The left ventricle was harvested and kept on dry ice. RIPA buffer (10 mM Tris-HCl pH 7.4, 1 mM EDTA, 0.5 mM EGTA, 1% NP-40, 0.1% sodium deoxycholate, 0.1% SDS, 140 mM NaCl) with protease and phosphatase inhibitors (Cell Signaling) lysed left ventricular heart tissue. The homogenates were placed on a rocker for 30 minutes at 4 °C, centrifuged at 14,000 g at 4 °C for 15 minutes, and the resulting supernatant was collected. Protein concentrations were measured using the DC™ protein assay (Bio-Rad). Protein extracts were separated by sodium dodecyl sulfate polyacrylamide gel electrophoresis on 4-20% gels. The gels were transferred to nitrocellulose membranes (Turbo blot, Bio-Rad) and blocked for 1 hour in 5% bovine serum albumin (BSA) in 1x TBS-T (10 mM Tris-HCl, pH 7.5, 150 mM NaCl, 0.1% Tween-20). Nitrocellulose membranes were incubated with primary antibodies diluted in blocking solution at 4 °C overnight. The primary antibodies included atrial natriuretic peptide (ANP, rabbit, 1:800 GeneTex), B-type natriuretic peptide (BNP, rabbit, 1:800 Abcam), type I collagen (Col1a1, rabbit, 1:1000 Novus Biologicals), type III collagen alpha 1 (Col3a1, rabbit, 1:1000 Novus Biologicals), hypoxia inducible factor 1 (HIF-1ɑ, rabbit, 1:750 Cell Signaling Technology), calcium/calmodulin dependent protein kinase II delta (CAMKII, rabbit, 1:800 GeneTex), phospho-inositol triphosphate receptor 1 and total-inositol triphosphate receptor 1 (ph-IP3R1, rabbit, 1:800 Cell Signaling Technology) (tot-IP3R1, rabbit, 1:1000 Cell Signaling Technology), cleaved caspase-3 (cleaved-Casp3, rabbit, 1:750 Cell Signaling Technology), LC3Β (LC3Β, rabbit, 1:750 Abcam), and SQSTM1 (p62, rabbit, 1:750 Abcam) . After incubation with primary antibodies, membranes were treated with horseradish peroxidase-conjugated species-specific secondary antibodies. Protein bands were detected using a chemiluminescent substrate (West Pico, ThermoFisher Scientific), imaged with ChemiDoc (BioRad), and band intensities were quantified using Image Lab software (BioRad). The total amount of protein loaded normalized data through visualization with Ponceau S solution (Sigma).

### Ex vivo assessment of cardiac electromechanical function (Dual Ca^2+^/V_m_ epicardial optical mapping)

Cytosolic Ca^2+^ and transmembrane potential (V_m_) were optically mapped from the epicardial surface of perfused hearts excised from a subset of mice in each group to determine if TNFR2 Ag prevented OA-induced changes in action potential duration (APD), cytosolic Ca2+ transient duration (CTD), and action potential conduction velocity. Mice were anesthetized with 5% isoflurane inhalation to provide a deep plane of anesthesia then placed supine with a nose-cone delivering 5% isoflurane to maintain a deep plane of anesthesia. The heart was then rapidly excised and placed in cold Krebs-Henseleit (KH) buffer solution supplemented with heparin. The aorta was cannulated and flushed with cold KH solution containing (in mM) 118 NaCl, 4.7 KCl, 1.25 CaCl2, 0.57 MgSO4, 1.17 KH 2PO4, 25 NaHCO3, and 6.0 glucose. The heart was then transferred to a perfusion system and retrograde perfused at a constant hydrostatic pressure of 75 mmHg with 37degC KH oxygenated with 95% O2 and 5% CO2. A pacing electrode was placed on the LV epicardium. Hearts were electromechanically uncoupled (blebbistatin) to prevent motion artifacts in the optical signals then stained with the fluorescent probes RH237 and Rhod2am to transduce action potentials and Ca2+ transients using 530nm excitation light. Action potentials and Ca2+ transients were simultaneously imaged using high speed CCD cameras during pacing at a cycle length of 145msec ^64, 65^. Optical mapping data were analyzed using custom MATLAB algorithms to measure average APD, CTD, and conduction velocity from the central LV epicardium^65^. APD was measured at 80% (APD80) to assess how OA impacts action potential repolarization and whether TNFR2 Ag prevents changes in APD80. CTD was also measured at 80% Ca2+ resequestration (CTD80) to assess how OA impacts Ca2+ resequestration and whether TNFR2 Ag prevents changes in CTD80. Conduction velocity was measured to assess whether action potential conduction was slower in OA mice and whether TNFR2 Ag prevents the slowing of conduction velocity.

## Statistical analysis

Data are shown using GraphPad Prism 10.0 software (San Diego, CA, USA) and are presented as mean ± SEM. Appropriate tests were used for each experiment. Two-way ANOVA (α = 0.05) was used for von Frey testing, subjects as factors and time as the repeated measure. One-way ANOVA analysis was used for multiple group comparison followed by Tukey post-hoc multiple comparison test. P-value <0.05 was considered significant.

## Results

### TNFR2 agonism attenuates DMM-induced joint mechanical hypersensitivity and cartilage degeneration in both sexes

To establish osteoarthritis in mice, we subjected male and female 12-week-old C57BL/6 mice to destabilization of the medial meniscus (DMM) surgery and evaluated mechanical sensitivity and joint pathology. Previous studies indicate the benefit of TNFR2 activation in alleviating chronic inflammatory pain ^31–37,40^. TNFR2 activation exerts immunosuppressive properties in rheumatoid arthritis and neuropathic pain models, therefore, we sought to ascertain if this effect extended towards pain associated with OA chronic inflammation ^31,37,48,49^. The TNFR2 agonist was administered intraperitoneally (IP) twice weekly at 1 mg/kg starting at 4 weeks post-surgery, following the onset of pain (Fig. 1A). Mice showed a progressive reduction in mechanical withdrawal thresholds beginning at 3 weeks post-surgery, indicating mechanical hypersensitivity. Both sexes developed significant hypersensitivity in the ipsilateral hind paw relative to contralateral controls. TNFR2 activation attenuated hypersensitization in both sexes (Fig. 1B-C)

**Figure 1.**
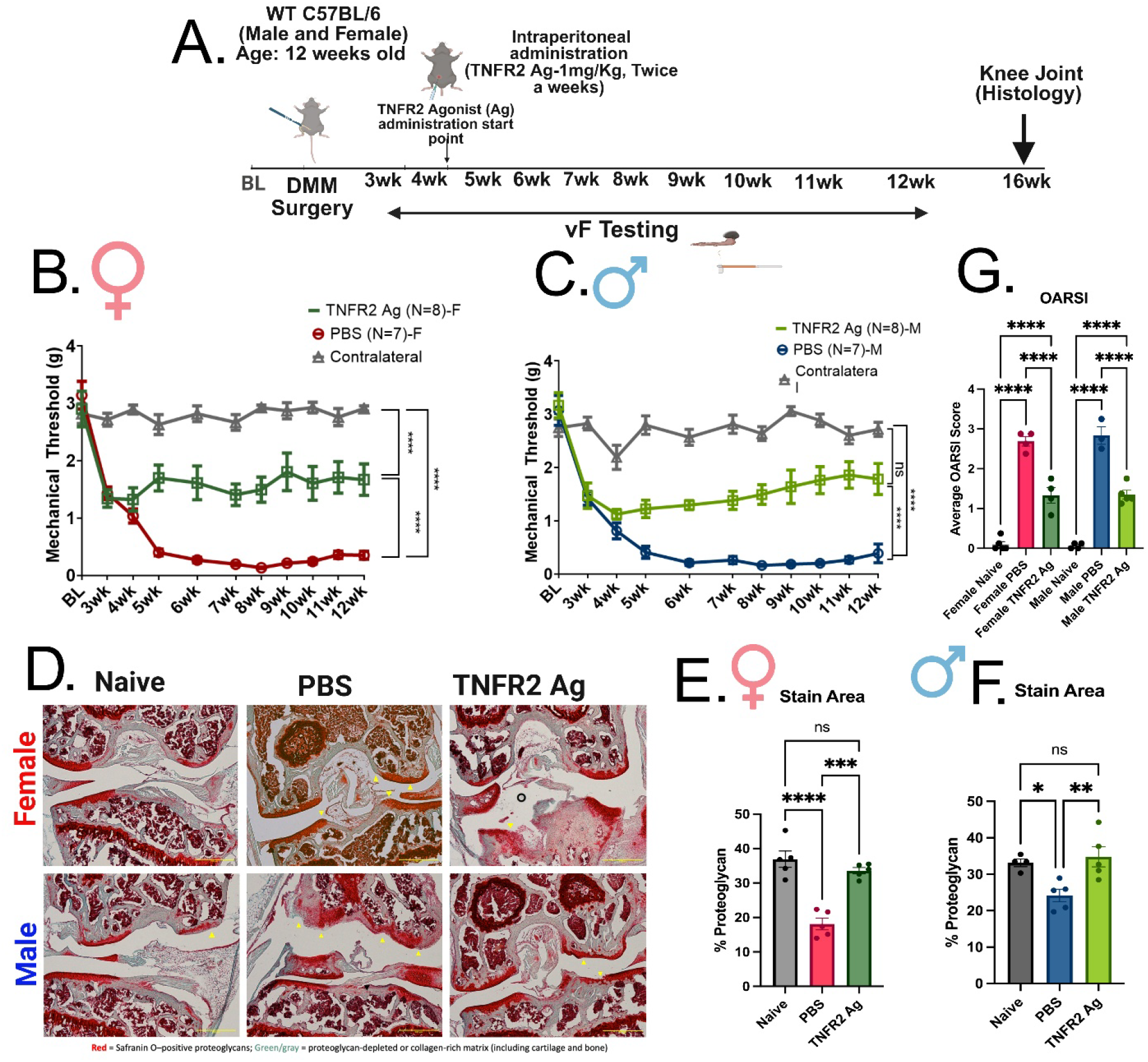
Destabilization of the medial meniscus induces pain and cartilage degeneration in the knee of mice. **(A)** Experimental timeline schematic. Twelve-week-old male and female *C57BL/6* mice underwent destabilization of the medial meniscus (DMM) surgery. Mechanical thresholds were assessed weekly from 3–12 weeks post-surgery using von Frey (vF) testing. Knee joint histology was performed at 16 weeks post-surgery. TNFR2 Agonist was provided twice weekly from 4 weeks onward. **(B-C)** Mechanical withdrawal thresholds over time in **(B)** female and **(C)** male mice. ipsilateral (surgery side) paws exhibited progressive and significant mechanical hypersensitivity compared to the contralateral side of the paw (control). Vehicle-treated mice exhibited progressive and significant mechanical hypersensitivity compared with TNFR2 agonist treatment. Data are mean ± S.E.M.; n = 7–8 per group. Two-way ANOVA with Sidak’s post-hoc test; ****p < 0.0001 **(D)** Representative Safranin-O–stained knee joint sections from naïve, vehicle-treated, and TNFR2 agonist-treated groups in female (top row) and male (bottom row) mice. OA groups show a marked reduction in proteoglycan staining (red) compared to naïve. Yellow arrows indicate surface fibrillation or damage. Scale bar: 200μm **(E-F)** Quantification of Safranin-O staining demonstrates significant proteoglycan loss in both female **(E)** and male **(F)** OA groups relative to naïve controls, with TNFR2 agonist-treated mice showing no change relative to naïve controls and significantly higher proteoglycan content relative to vehicle-treated animals. Data are mean ± S.E.M.; n = 4–5 per group. One-way ANOVA with Tukey’s post-hoc test; * p < 0.05, **p < 0.01, ***p < 0.001, and ****p < 0.0001. **(G)** Quantification of cartilage degeneration using OARSI scoring with female and male OA groups exhibiting significantly higher scores compared to naïve controls. TNFR2-agonist treated mice had higher scores relative to naïve controls but reduction compared to vehicle-treated animals.

Histological analyses of knee joints at 16-weeks post-surgery provided clear evidence of cartilage damage in PBS-treated animals. Safranin-O staining, labeling proteoglycan content in articular cartilage, demonstrated substantial proteoglycan depletion in DMM joints of PBS-treated mice compared to naïve controls at the tibial plateau. TNFR2 activation restored proteoglycan content (Fig. 1D). Quantitative analysis revealed significant reductions in Safranin-O–positive regions across both sexes in PBS-treated mice compared to naïve controls, indicating widespread proteoglycan loss, a hallmark of OA pathology. Therapeutic TNFR2 agonist administration rescued proteoglycan loss (Fig. E-F). Loss of Safranin-O staining intensity corresponded with areas of surface fibrillation and structural disruption, indicating progressive cartilage degeneration. Consistent with these findings, OARSI scoring further substantiated cartilage degeneration and structural damage in PBS-treated mice. Compared to naïve controls, DMM-operated joints exhibited elevated OARSI scores reflecting the combined severity of proteoglycan depletion, cartilage surface irregularity, and structural breakdown. While the OARSI score of TNFR2 agonist-treated mice remained significantly elevated compared to naive controls, it attenuated the overall severity of OARSI scores compared to mice treated with PBS (Fig. 1G). Importantly, this scoring method provided a semiquantitative assessment paralleling histological observations, further validating the severity of the OA phenotype induced by DMM, while highlighting therapeutic attenuation via TNFR2 activation.

### DMM surgery causes sex-specific cardiac dysfunction, while prophylactic TNFR2 activation prevents female diastolic dysfunction and delays the onset of male systolic dysfunction

#### Female OA and Diastolic dysfunction

Inflammation mediates the progression of heart failure (HF)^50^. Epidemiological evaluation of patient populations indicates a strong correlation between osteoarthritis and CVD ^51^. Given the potential for confounding factors, including weight gain, in epidemiological studies, we used echocardiography to assess DMM mice of both sexes biweekly while ensuring consistent weights between groups. Mice’s weights did not exceed the expected weights of Jackson C57 mice of comparable age (Fig. S1 A). Female mice developed diastolic dysfunction with disease onset at 8 weeks post-DMM surgery. They showed a significantly altered E/A ratio (initial decrease then increase), elevated E/e’ ratio, prolonged intraventricular relaxation time (IVRT), and increased thickness of the posterior wall at diastole without changes to ejection fraction when compared to baseline control measurements from 8 weeks to the experimental endpoint, 16 weeks (Fig. 2C-G). Longitudinal strain analysis demonstrated an increased global longitudinal strain, decreased global radial strain, and increased outer wall delay (Fig. S1 C-G). Mean arterial pressure (MAP) was assessed longitudinally throughout the experiment, analyzing potential blood pressure changes. At 6 weeks, vehicle-treated mice had a significant elevation of MAP compared to baseline and TNFR2 agonist-treated mice. Elevated MAP values in vehicle-treated mice persisted until the experimental endpoint (Fig. 2H). Morphometric analysis at 16 weeks revealed increased heart weight-to-tibia length ratio (HW/TL³) (Fig. 2I), consistent with concentric hypertrophy. TNFR2 agonism (1 mg/kg) following onset of pain, 4 weeks post-surgery in female OA mice, prevented diastolic dysfunction. Hematoxylin and eosin (H&E) staining of female heart sections at 16 weeks showed reduced LV chamber diameter and increased wall thickness, consistent with concentric hypertrophy (Fig. S2). Masson trichrome staining (MTS) corroborated this, showing increased cartilage deposition in vehicle-treated female mice (Fig. 2J-K). Also, we performed optical mapping at the end of the experiment. We found that Conduction velocity was significantly reduced in vehicle-treated mice, but the TNFR2 agonist prevented this change, showing a preservation of electromechanical function (Fig 2L). Assessment of female cardiac electromechanical function revealed slower conduction velocity with no change in action potential and calcium transient durations, consistent with increased fibrosis that disrupts electrical coupling (Fig. 2 M-N). Together, these findings indicate impaired relaxation, hypertrophy, and progressive diastolic dysfunction in females.

**Figure 2.**
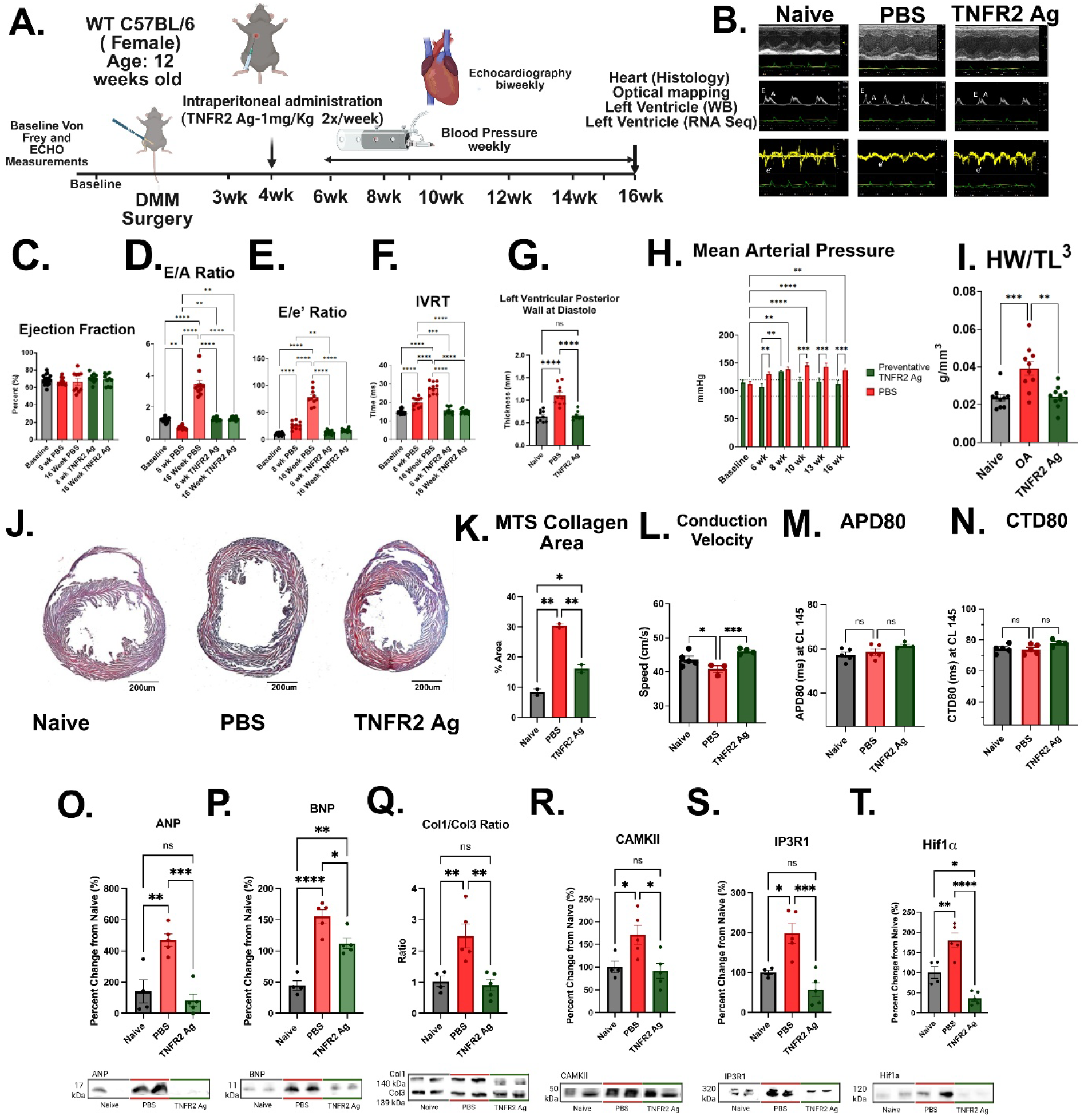
Osteoarthritis (OA) induces diastolic dysfunction in female mice. **(A)** Timeline showing heart assessment and TNFR2 activation. (Preventive systemic administration of 1 mg/kg TNFR2 agonist). **(B)** Echocardiographic tracings. Representative M-mode, pulse-wave Doppler, and tissue Doppler recordings from naïve, vehicle-treated, and TNFR2 agonist-treated mice illustrate preserved systolic function with impaired diastolic relaxation, prevented by prophylactic administration of the TNFR2 agonist. **(C-G) Female cardiac function.** Quantitative analyses showed no change in ejection fraction at 8- and 16-weeks post-OA compared to baseline **(C),** but PBS-treated mice developed progressive diastolic dysfunction characterized by changes in E/A **(D)** and E/e′ ratios **(E),** IVRT **(F),** while agonist-treated mice showed no difference to baseline control. **(G)** Significantly increased posterior wall thickness at diastole indicated relaxation abnormalities. TNFR2 Ag attenuated it. **(H) Longitudinal mean arterial pressure** in female vehicle-treated mice increased starting at 6 weeks and remained elevated throughout the experiment. TNFR2 agonist-treated mice remained in the normal range. **(I) Morphometric assessment of hypertrophy.** Female vehicle-treated mice at 16 weeks exhibited a significant increase in heart weight-to-tibia length ratio (HW/TL³) when compared to both naïve controls and TNFR2 agonist treatment. **(J)** Masson Trichrome Stain and Analysis. Representative stains of female naive, vehicle-treated, and TNFR2 agonist-treated mice showed increased collagen deposition (blue) among cardiomyocytes (red) in vehicle-treated mice. Scale bar: 200μm **(K)** Quantitative analysis showed a significant elevation of total collagen in vehicle-treated mice. TNFR2 agonist-treated mice showed a reduction relative to vehicle-treated mice, but levels remained elevated compared with naive hearts. **(L) Cardiac electrical signaling.** Female vehicle-treated mice showed a significant reduction in conduction velocity compared to naive and TNFR2 agonist-treated mice. **(M-N) Electromechanical function.** Female mice showed no differences in action potential durations (APD) **(M)** and calcium transient durations (CTD) **(N)** across groups. **(O-T) Western blot analysis of cardiac biomarkers.** Female OA vehicle-treated mice show significant increases in **ANP (O), BNP (P), Col1/Col3 ratio (Q), CAMKII (R), IP3R1 (S),** and **HIF1ɑ (T),** indicative of cardiac stress, oxidative stress, and fibrosis. TNFR2 agonist -treatment decreased ANP, BNP, Col1/Col3 ratio, CAMKII, IP3R1, and HIF1ɑ when compared to vehicle-treated (PBS) mice. Data presented as mean ± S.E.M.; statistical analysis by one-way ANOVA with post hoc multiple comparisons. *P < 0.05, **P < 0.01, ***P < 0.001. Group sizes: n=8–10 for echocardiographic measures; n=5 for morphometry.

Cardiac biomarkers were assessed in left ventricular tissue through Western blot^58–59^. Female vehicle-treated mice showed a significant increase in ANP, BNP, Col1/Col3, CAMKII, IP3R1, and Hif1ɑ at 16 weeks post-surgery, indicative of cardiac stress, fibrotic remodeling, and oxidative stress (Fig. 2 O-T). TNFR2 agonist-treated and control mice showed no differences in ANP, Col1/Col3 ratios, CAMKII, IP3R1, and Hif1ɑ. In BNP, TNFR2 agonist-treated mice remained elevated relative to naive controls, while vehicle-treated mice were significantly reduced (Fig O-T). TNFR2 agoinst administration, starting at 4 weeks post-surgery, prevented diastolic dysfunction in females when administered prophylactically.

### Male OA and Systolic Dysfunction

Male mice developed systolic dysfunction with disease onset between 10-12 weeks post-surgery. Mice’s weights did not exceed the expected weights of Jackson C57 mice of comparable age (Fig. S1 B). Male OA mice exhibited a progressive decline in systolic function. Echocardiography conducted biweekly showed reduced contractility relative to both naïve males and baseline (Fig 3A). They showed decreased ejection fraction, elevated E/e’ ratio, increased intraventricular contraction time, decreased thickness of the posterior wall at systole, and, when compared to controls and baseline measurements (Fig. 3C-G). Longitudinal strain analysis of male mice showed an increased global longitudinal strain (GLS), decreased global radial strain (GRS), and increased outer wall delay (OWD) (Fig. S1 B-H). H&E staining of male heart sections at 16 weeks post-surgery showed no differences in chamber diameter and wall thickness when compared to controls (Fig S2). (Fig 3H, S3) Morphometric assessment showed no increase in HW/TL³ at 16 weeks (S3E) but a significant increase at 20 weeks (Fig 3H). Mean arterial pressure (MAP) assessment showed a progressive decrease relative to baseline throughout the experiment (Fig. 3I). (Fig J-K) Assessment of male cardiac electromechanical function showed no change in conduction velocity but a decreased action potential duration, potentially suggesting increased adrenergic activation in male hearts. These findings indicate systolic dysfunction in the males contrasting with the female’s diastolic dysfunction. Overall, this suggests a divergence in male and female hearts’ response to OA-induced chronic inflammation.

**Figure 3.**
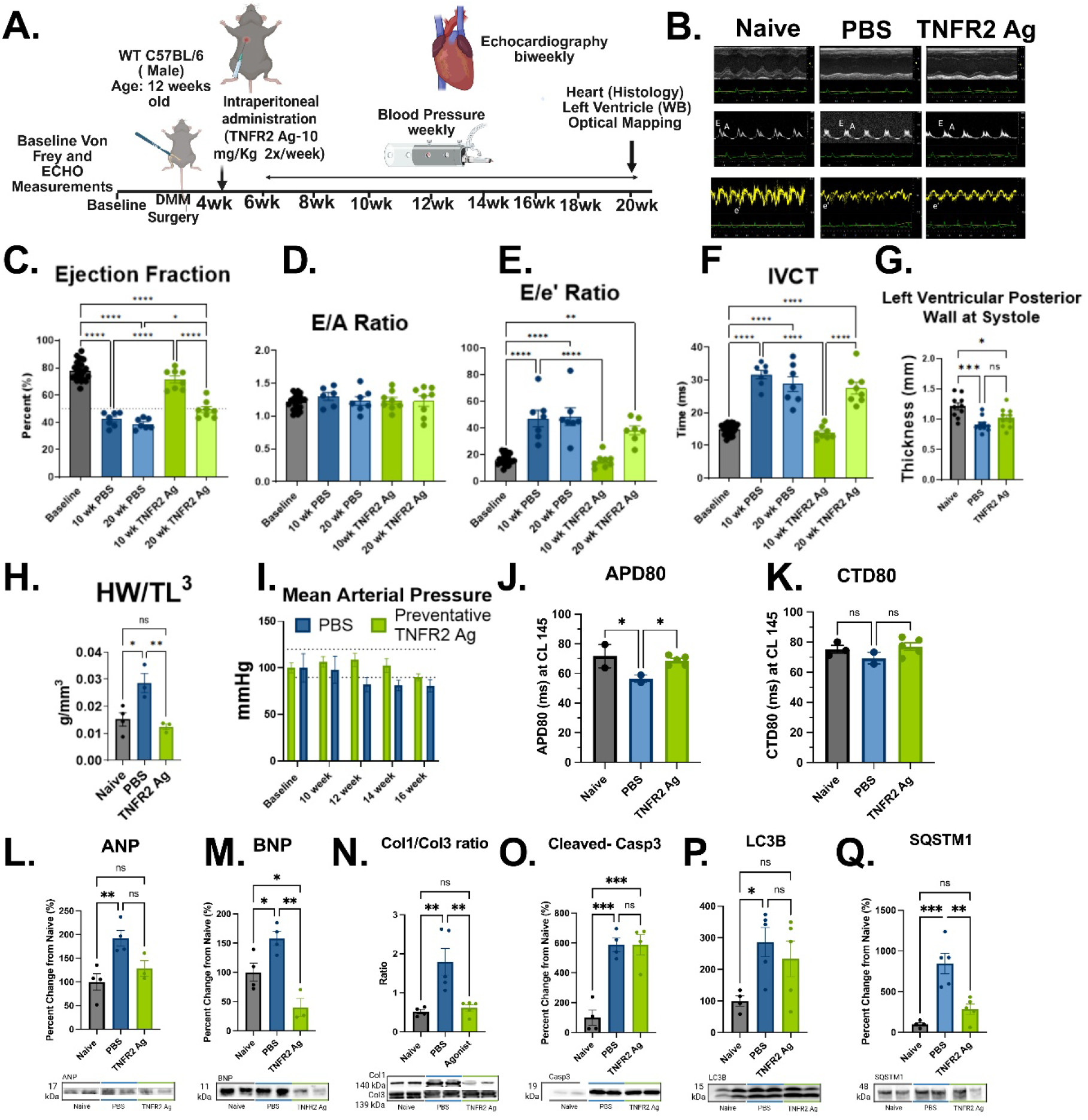
Osteoarthritis (OA) induces systolic dysfunction in male mice. **(A)** Timeline showing heart assessment and TNFR2 activation (preventive systemic administration of TNFR2 agonist (10 mg/kg). **(B)** Echocardiographic tracings. Representative M-mode, pulse-wave Doppler (E/A ratio), and tissue Doppler (E/e′) recordings from naïve, vehicle-treated, and TNFR2 agonist-treated illustrate impaired systolic function, improved by prophylactic administration of the TNFR2 agonist. **(C-F)** Male cardiac function (onset of systolic dysfunction starts at 10 weeks after DMM surgery). Quantitative analyses showed males maintained the E/A ratio at 10- and 20-week post-OA when compared to baseline **(D)**, but developed progressive systolic dysfunction characterized by decreased ejection fraction **(C)** and increased E/e′ ratios **(E),** while the agonist-treated mice showed no difference from naïve controls at 10 weeks, and only a moderate increase from vehicle-treated mice at 20 weeks. Similarly, increased IVCT in vehicle-treated mice **(F)**; however, the agonist mitigated the impaired contraction. **(G)** Decreased thickness of the posterior wall at systole in vehicle-treated mice, indicating contraction abnormalities. **(H)** Morphometric assessment of hypertrophy. Male vehicle-treated mice exhibited a significant increase in heart weight-to-tibia length ratio (HW/TL³) when compared to both naïve and TNFR2 agonist-treated mice. **(I)** Longitudinal mean arterial pressure. Male vehicle-treated mice showed decreased mean arterial pressure starting at 10 weeks and continuing throughout the experiment. **(J-K)** Electromechanical Function. Male vehicle-treated mice showed significant reductions in action potential duration **(J)** with no change in calcium transients **(K)**, indicating altered electrical signaling. **(L-Q)** Western blot analysis of cardiac biomarkers. Male OA vehicle-treated mice show significant increases in ANP **(L)**, BNP **(M)**, Col1/Col3 ratio **(N)**, cleaved-caspase 3 **(O)**, LC3β **(P)**, and SQSTM1 **(Q)**, indicative of cardiac stress, fibrosis, apoptotic, and autophagic responses. TNFR2 agonist treatment decreased ANP, Col1/Col3 ratio, BNP, and SQSTM1 compared with vehicle-treated mice. Data presented as mean ± S.E.M.; statistical analysis by one-way ANOVA with post hoc multiple comparisons. *P < 0.05, **P < 0.01, ***P < 0.001. Group sizes: n=8–10 for echocardiographic measures, n=5 for morphometry.

Biochemical analysis of ANP, BNP, Col1/Col3 ratio, cleaved caspase-3, LC3β, and SQSTM1 in left ventricular tissue showed significant increases at 20 weeks in vehicle-treated mice, indicating cardiac stress, autophagic, and apoptotic pathways. Prophylactic TNFR2 agonist treatment at 10 mg/kg significantly reduced BNP, Col1/Col3, and SQSTM1 levels compared with vehicle-treated mice, but failed to reduce cleaved caspase-3 and LC3β (Fig 3L-Q). Overall, these proteins provide evidence for eccentric remodeling in the male mice.

TNFR2 agonist was administered at 10 mg/kg given the ineffectiveness of a 1 mg/kg dose in preventive groups. Furthermore, the preventive low-dose TNFR2 agonist (1 mg/kg) administered from 4 weeks onward showed no significant change compared with vehicle-treated mice (Fig S3). In contrast, the preventive high dose of the TNFR2 agonist (10 mg/kg) prevented the onset of systolic dysfunction at 10 weeks (Fig 3). However, efficacy declined over time, as indicated by a progressive decrease in ejection fraction despite improvement relative to vehicle-treated mice.

### Left ventricular cardiovascular gene expression following DMM is sexually dimorphic

To further elucidate the mechanisms of CVD and the underlying sexual dimorphism in DMM mice, we performed left ventricular bulk RNA sequencing. PCA analysis revealed distinct clustering profiles between male and female diseased groups, highlighting divergent disease phenotypes (Fig. 4A). Female mice had 531 differentially expressed genes, and male mice had 204, with only 19 genes shared between groups (Fig. 4D).

**Figure 4.**
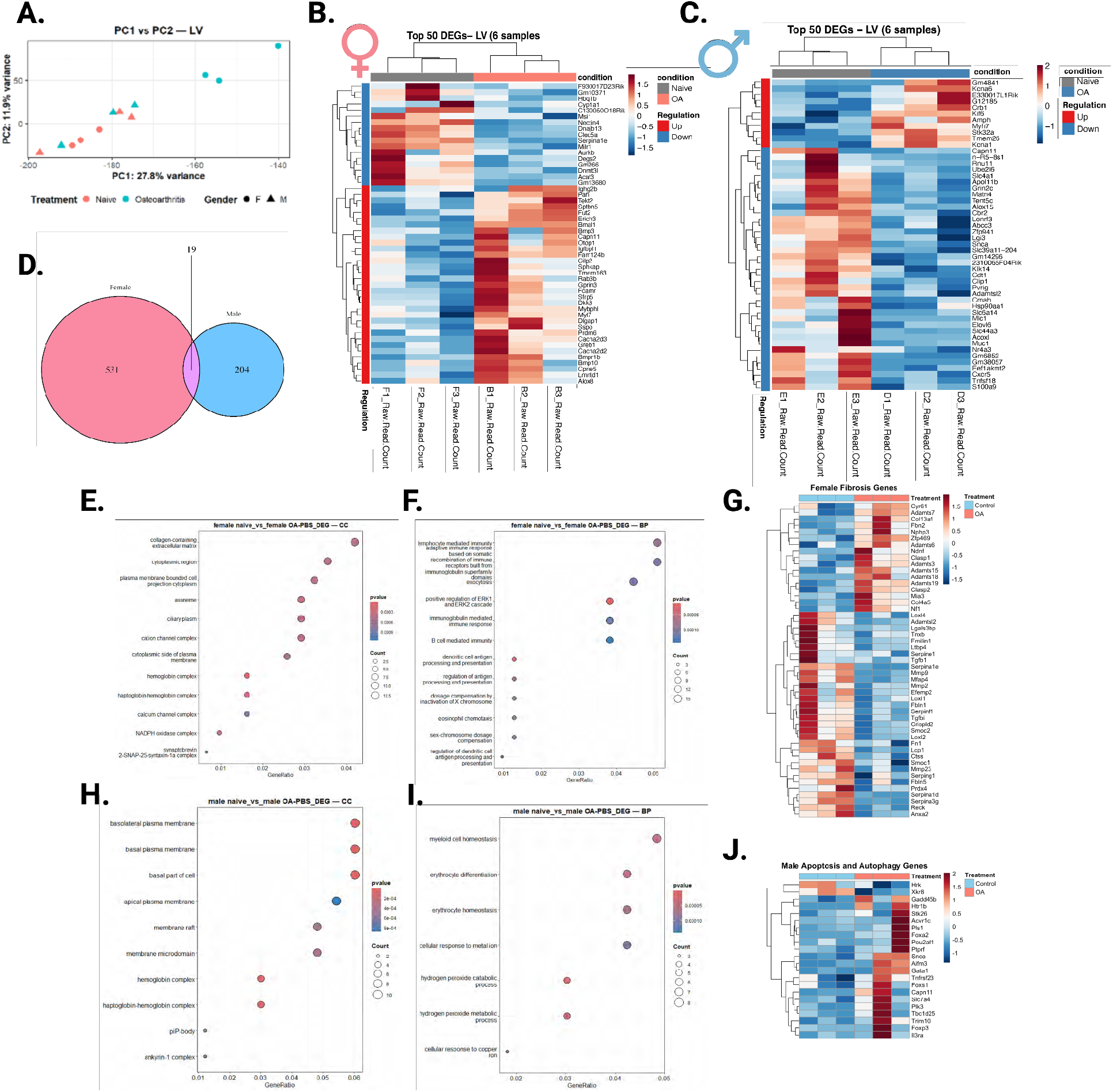
Transcriptomic analysis of left ventricular tissue reveals distinct disease pathologies in males and females following novel OA-induced heart failure model. **(A) Principal Component Analysis.** Principal component analysis (PCA) of normalized RNA-sequencing data assessed overall transcriptional variation in left ventricular tissue from naïve and OA mice across both sexes. **(B-C) Comparison of Male and Female Gene Expression.** Heatmaps illustrate the expression of the top 50 genes in left ventricular (LV) tissue, highlighting pathways related to cytoskeletal remodeling, immune regulation, and inflammatory signaling in female (B) and male (C) mice. **(D)**The Venn diagram shows the distribution of differentially expressed genes (DEGs) in left ventricular tissue between Males and Females . **(E-F) Female pathway analysis.** Gene Ontology enrichment analysis of differentially expressed genes in female mice following OA-induced diastolic dysfunction, organized by Cellular Component (CC) (E) and Biological Process (BP) (F). **(G)** Heatmap of expressed genes in GO:0030198 related to extracellular matrix organization and fibrosis. Samples were clustered based on normalized expression values. **(H-I) Male pathway analysis.** Gene Ontology enrichment analysis of differentially expressed genes in male mice following OA-induced systolic dysfunction, organized by Cellular Component (CC) (H) and Biological Process (BP) (I). **(J)** Heatmap of expressed genes in GO:0097194, GO:1904716, and GO:0010507 related to apoptosis and autophagy. Samples were clustered based on normalized expression values.

Cellular Component gene ontology pathway analysis in female mice showed differences in ion channel and oxygen handling gene expression. Calcium-associated and collagen-associated genes were differentially expressed, a hallmark of diastolic dysfunction and fibrotic remodeling (Fig. 4E). Biological process gene ontology pathway analysis revealed differences in B cell activity and immune responses (Fig. 4F). Meanwhile, cellular component pathway analysis of male mice showed changes to plasma membrane structure while biological processes pathway analysis emphasized homeostatic processes and ion channel regulation (Fig. 4H-I). Overall, the diverging gene expression patterns in males and females support the diverging cardiovascular response seen through functional measurements.

Upregulated genes in female OA hearts revealed enrichment of pathways related to cardiac remodeling *(Myl7, Bmp10*), fibrotic signaling (*Bmpr1b, Sfrp5*), calcium handling (*Cacna2d2/3*), circadian/metabolic regulation (*Bmal1*), and inflammatory and immune response genes (*Ighg2b, Fcamr, Clec5a*), all mechanistically linked to diastolic dysfunction pathophysiology ^52–57^. Females showed downregulation of specific genes involved in metabolism (*Acat3, Degs2*) (Fig. 4B) ^12^. Male diseased hearts showed upregulation of hypertrophic (*Myh7*) and contractile genes (*Kcna1, Kcna6*), alongside downregulation of metabolism associated genes (*Mlc1, Acoxl, Elovl6*) (Fig. 4C) ^58–62^. Further analysis indicated strong differential expression of fibrotic pathway genes in female mice and apoptotic and autophagic genes in male mice (Fig. 4G, J). These gene signatures correlate with the sexually dimorphic dysfunction observed by echocardiography, indicating a sex-specific CVD pattern.

### Therapeutic TNFR2 activation starting at the onset of disease treated diastolic changes in female mice and improved systolic changes in male mice

Given clinical paradigms, we investigated the potential therapeutic effects of the TNFR2 agonist at the onset of cardiovascular dysfunction. Female mice showed a significant change from baseline values at 8 weeks before beginning TNFR2 agonist- (1 mg/kg) or vehicle-treatment (Fig. 5 C-E). After 2 weeks of TNFR2 agonist administration, diastolic parameters (E/A ratio and E/e’) exhibited marked improvement from vehicle-treated mice (Fig. F-I). At 16 weeks, therapeutic administration of the TNFR2 agonist significantly improved conduction velocity (Fig 5 J). The TNFR2 agonist therapeutic effects persisted to the end timepoint of 16 weeks for the female mice. Western blot of ANP, BNP, Col1/Col3. Vehicle-treated mice showed elevated ANP (Fig. 5M), BNP (Fig. 5N), and Col1/Col3 ratio (Fig. 5O) compared to both naive and TNFR2 agonist-treated mice.

**Figure 5.**
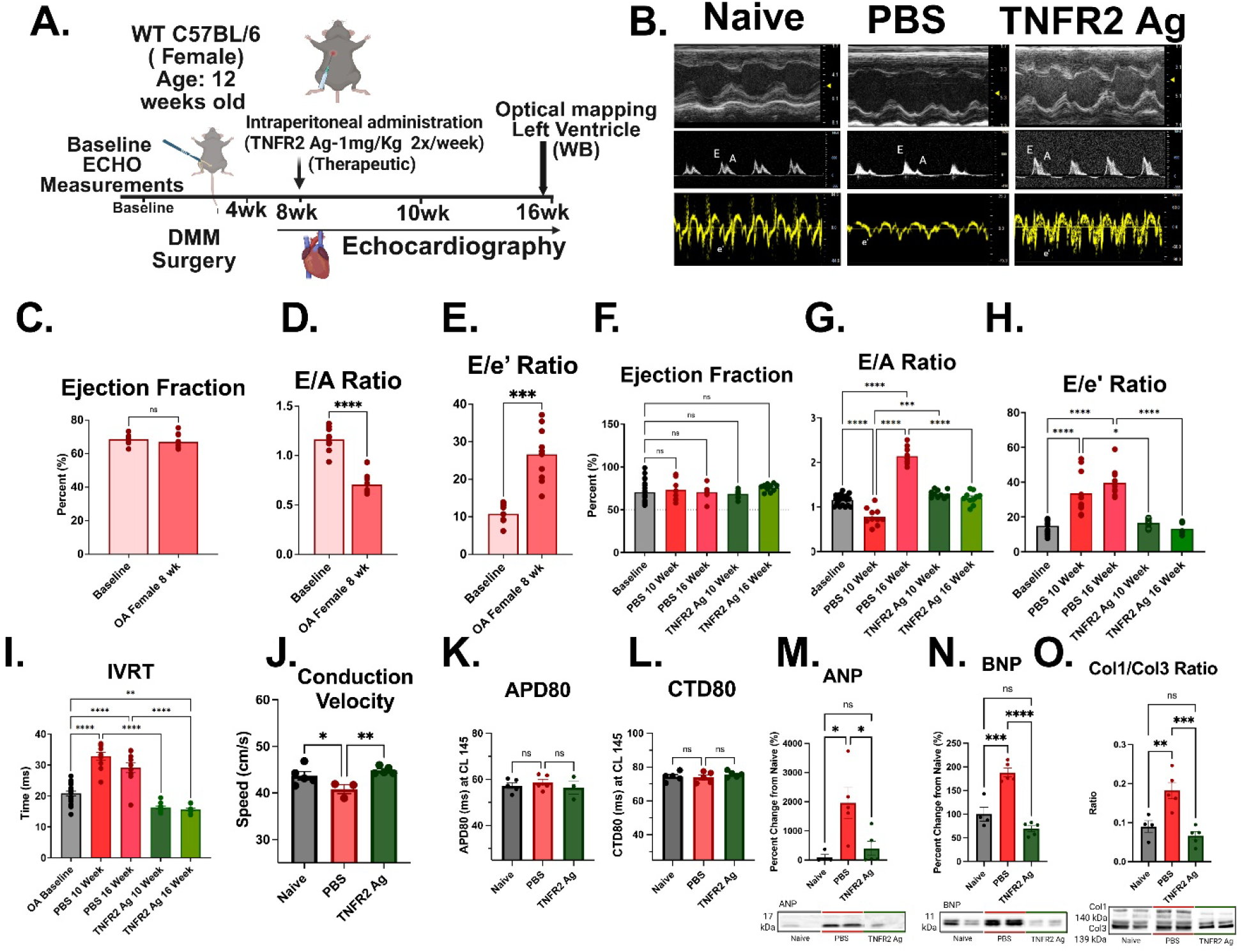
The therapeutic TNFR2 agonist successfully rescued Osteoarthritis (OA)-induced diastolic dysfunction in female mice. **(A).** Timeline showing heart assessment and therapeutic administration of TNFR2 agonist. **(B) Echocardiographic tracings.** Representative M-mode, pulse-wave Doppler (E/A ratio), and tissue Doppler (E/e′) recordings from naïve, vehicle-treated, and TNFR2 agonist-treated mice illustrate impaired diastolic function, with TNFR2 agonist treatment rescuing the phenotype. (**C-E) Induction of Cardiac Dysfunction.** Quantitative analysis shows the onset of cardiac dysfunction as assessed by an unchanged ejection fraction (C), decreased E/A ratio (D), and elevated E/e’ ratio (E) at 8 weeks following DMM surgery in female mice. Administration of the TNFR2 agonist began at 8 weeks, a therapeutic time point for female mice. **(F-I) Female cardiac function.** Quantitative analyses showed vehicle-treated females maintained ejection fraction at 10-and 16-weeks post-OA compared to baseline (F) but developed progressive diastolic dysfunction characterized by increases in E/A (G), E/e′ ratios (H), and IVRT (I). TNFR2 agonism restored parameters to baseline levels 2 weeks after the first injection and maintained them through the 16-week endpoint. **(J-L) Electromechanical Function.** Vehicle-treated mice showed a significant reduction in conduction velocity, which TNFR2 agonism rescued (J), with no changes in action potential duration (K) and calcium transients (L) **(M-O). Western blot of ANP, BNP, Col1/Col3.** Vehicle-treated mice showed elevated ANP (M), BNP (N), and Col1/Col3 (O) ratio compared to both naive and TNFR2 agonist-treated mice. Data presented as mean ± S.E.M.; statistical analysis by one-way ANOVA with post hoc multiple comparisons. *P < 0.05, **P < 0.01, ***P < 0.001. Group sizes: n=8–10 for echocardiographic measures; n=5 for morphometry.

Male mice developed a decreased ejection fraction and increased E/e’ at 10 weeks (Fig. 6C-E). TNFR2 agonist-treated mice showed significant improvements in ejection fraction (Fig 6F), E/e’ (Fig 6H), and IVCT (Fig 6I) 2 weeks after the first dose. However, parameters were still reduced compared to naive control and showed patterns more similar to the vehicle-treated mice at later time points (Fig. 6F-I). Therapeutic TNFR2 agonist administration failed to affect action potential duration compared with vehicle-treated controls (Fig. 6J). However, there is no change in conduction velocity (Fig 6K). Further studies investigating TNFR2 agonist dosage and effectiveness in male mice are warranted to elucidate the role of TNFR2 signaling in systolic dysfunction. In Western blot analysis. Vehicle-treated males showed elevations in ANP (Fig. 6L), BNP (Fig. 6M), and the Col1/Col3 ratio (Fig. 6N) with TNFR2 agonist treatment; only the Col1/Col3 ratio improved significantly.

**Figure 6.**
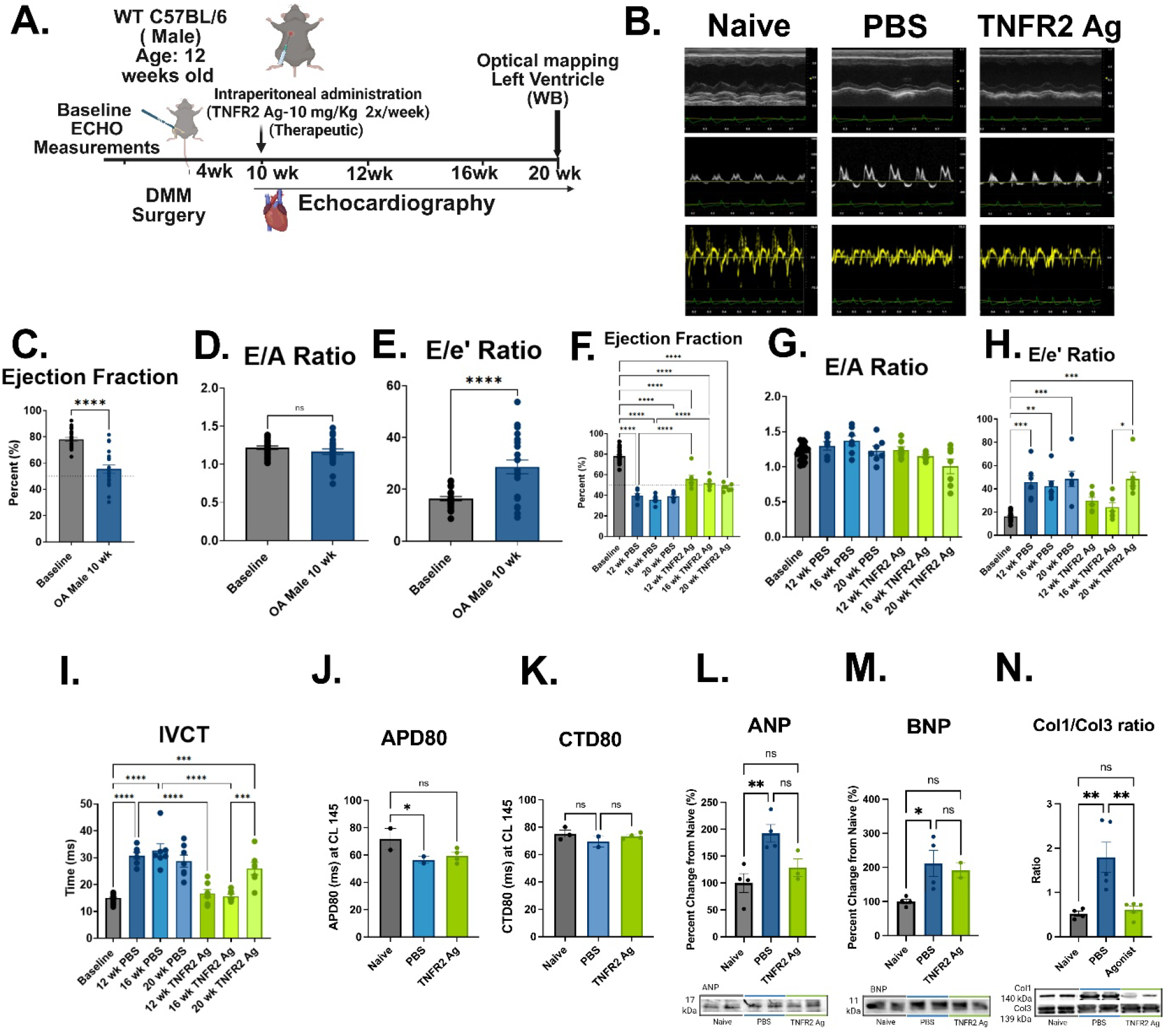
The therapeutic TNFR2 agonist administration improved OA-induced systolic dysfunction in male mice. **(A)** Experimental timeline of therapeutic administration of TNFR2 agonist. **(B) Echocardiographic tracings.** Representative M-mode, pulse-wave Doppler (E/A ratio), and tissue Doppler (E/e′) recordings from naïve, vehicle-treated, and TNFR2 agonist-treated mice illustrate that impaired systolic function remains unchanged with TNFR2 agonist treatment (C-E). **Induction of Cardiac Dysfunction.** Quantitative analysis shows the onset of cardiac dysfunction, with decreased ejection fraction (C), unchanged E/A ratio (D), and elevated E/e’ ratio (E) at 10 weeks following DMM surgery in male mice. Administration of the TNFR2 agonist began at 10 weeks, a therapeutic time point for male mice. **(G-I) Male cardiac function.** Quantitative analyses showed vehicle-treated males maintained the E/A ratio at 12- and 20-weeks post-OA compared to baseline (G), but developed progressive systolic dysfunction characterized by a reduction in ejection fraction (F) and an increased E/e′ ratio (H). TNFR2 agonist treatment showed a moderate improvement at 12 weeks compared with vehicle-treated mice, but remained significantly different from controls and baseline measurements. At 20 weeks after DMM surgery, TNFR2 agonist-treated mice show no statistically significant difference from vehicle-treated mice. **(J-K) Electromechanical Function.** Male vehicle-treated mice showed decreased action potential duration (J), with TNFR2 agonism not showing a significant improvement. No change in conduction velocity (K). **(L-N) Western blot analysis.** Vehicle-treated males showed elevations in ANP (L), BNP (M), and the Col1/Col3 ratio (N), with TNFR2 agonist treatment significantly improving only the Col1/Col3 ratio. Data presented as mean ± S.E.M.; statistical analysis by one-way ANOVA with post hoc multiple comparisons. *P < 0.05, **P < 0.01, ***P < 0.001. Group sizes: n=8–10 for echocardiographic measures; n=5 for morphometry.

## Discussion

### Osteoarthritis induces cardiovascular dysfunction

The present study indicates osteoarthritis (OA) directly induces cardiac dysfunction, independent of the cardiovascular risk factors of physical inactivity and obesity. Using the destabilization of the medial meniscus (DMM) model in C57BL/6J mice, we established a novel preclinical model in which OA drives cardiac dysfunction. Epidemiological studies consistently report a 3-fold increased incidence of CVD in OA patients^4–8^. However, the causal nature of this association remained unclear due to confounding variables: mobility, weight, and nonsteroidal anti-inflammatory drug use. We addressed this limitation, controlling body weight across groups, utilizing a surgical OA model without dietary manipulation or immobilization. Our data confirms OA alone precipitates cardiac dysfunction in the absence of confounders, reframing OA as an independent driver of cardiac remodeling and dysfunction rather than a CVD-associated comorbidity. Overall, we demonstrate that OA is associated with progressive CVD *in vivo*, supporting the relevance of systemic inflammation to CVD. Further studies integrating clinical comorbidities, such as obesity, metabolic syndrome, and physical inactivity, with OA are needed to delineate likely additive contributions to the clinical phenotype.

### Osteoarthritis induces sexually dimorphic CVD

Male and female mice show dimorphic cardiovascular phenotypes: males develop systolic dysfunction, and females develop diastolic dysfunction. Abnormal E/A ratios, E/e’ ratios, prolonged IVRT, and unchanged ejection fraction from 8-16 weeks post-DMM surgery indicated diastolic dysfunction in female mice. In contrast, male mice exhibited a progressive decline in ejection fraction beginning at 10 weeks with prolonged IVCT and elevated E/e’ ratios. Elevated atrial natriuretic peptide (ANP), B-type natriuretic peptide (BNP), and collagen I/III ratios in both sexes corroborated functional findings at the molecular level. Female mice showed measurably increased hypertrophy and collagen deposition, which, when paired with decreased conduction velocity and unchanged calcium transient durations, support fibrosis-driven diastolic impairment while cardiomyocytes’ contractile function remains preserved. Meanwhile, male mice developed a phenotype more consistent with eccentric remodeling with increased cell death corresponding to stress-pathway (adrenergic) induced reduction action potential duration alongside the systolic impairment. The sex-specific cardiac phenotypes closely mirror clinical cardiac HF epidemiology. HFpEF disproportionately affects women, particularly post menopause, whereas HFrEF predominates in men^16,18^. This dual-phenotype model recapitulates the heterogeneity of human HF within a single, inflammation-driven system and provides a platform for investigating sex-specific therapeutic responses ^17,18^.

Bulk RNA sequencing of left ventricular tissue further demonstrated distinct molecular changes in these cardiac phenotypes. Principal component analysis demonstrated clear separation between sex-specific disease groups with only 19 differentially expressed genes overlapping. Females showed upregulation of fibrotic signaling, calcium handling genes, cardiac remodeling regulators, B cell-mediated immunity, inflammatory signaling and circadian/metabolic mediators, consistent with other preclinical HFpEF models. These results support the hypothesis that immune activation propagates from arthritic joints to the myocardium, driving fibrotic and diastolic phenotype. In males, upregulation of hypertrophic and potassium channel genes, alongside downregulation of metabolic genes, aligns with the contractile impairment and systolic dysfunction observed in HFrEF models. Overall, the transcriptomic profiles establish distinct molecular signatures for OA-induced diastolic and systolic dysfunction, identifying candidate pathways for sex-specific therapeutic targeting. Despite a common upstream inflammatory stimulus, male and female mice exhibit divergent functional and molecular remodeling, indicating sex influences downstream disease expression. These findings argue against pooled interpretation and suggest mechanism-based therapies require evaluation within sex-specific biological contexts.

### Prophylactic TNFR2 agonism prevents female and delays male CVD

The therapeutic rationale for selective TNFR2 agonism derives from the divergent roles of TNF receptors in cardiac pathophysiology. TNFR1 activation promotes proinflammatory signaling, apoptosis, and pathological remodeling, whereas TNFR2 signaling exerts is cardioprotective effects by supporting regulatory T cell expansion, mitochondrial fusion and suppression of NF-*κ*B-mediated inflammation. Prior clinical trials targeting TNFα nonselectively in HF yielded few results, likely because global TNF inhibition abrogates beneficial TNFR2 signaling alongside pathological TNFR1 signaling^24,25, 28–30^. Genetic ablation studies reinforce this paradigm: TNFR1 knockout exacerbates remodeling, fibrosis, and mortality^24,25,29,30^. Selective TNFR2 agonism therefore represents a strategy to harness the protective arm of TNF signaling without the deleterious effects of TNFR1 activation^31,32,33,37,63^.

In our study, preventive TNFR2 agonist administration (1 mg/kg) in female mice beginning 4 weeks after DMM-surgery prevented diastolic impairment with echocardiographic measurements remaining at baseline levels through 16 weeks, mean arterial pressure unchanged, and cardiac biomarkers comparable to controls. In contrast, in males (preventive dose of 1 mg/kg), the agonist failed to prevent systolic decline, potentially reflecting sexually divergent cardiovascular phenotypes. Prophylactic administration at 10 mg/kg TNFR2 delayed systolic dysfunction, with efficacy decreasing over time, as ejection fraction progressively declined. When initiated before dysfunction, the intervention prevented and delayed dysfunction in females and males, respectively, supporting targeted early disease approaches. The persistence of molecular abnormalities suggests treatment modifies but may not fully attenuate pathogenicity. This data supports further evaluation of dosage and comparison to current therapies.

### Therapeutic TNFR2 agonism reverses female and slows male CVD

In females, administration of a TNFR2 agonist (1 mg/kg) at the onset of established diastolic dysfunction (8 weeks post-DMM) reversed echocardiographic changes within 2 weeks, with improvements persisting through the 16-week endpoint. Therapeutic administration in males (10 mg/kg) following established systolic dysfunction produced initial improvements in echocardiographic indices, but the benefits did not persist at later time points. Male TNFR2 agonist-treated mice remained improved compared to vehicle-treated mice but were significantly worse than controls. This dose-dependent, time-limited efficacy in males suggests that TNFR2 agonism modulates, but does not override, the pathological cascade driving systolic dysfunction. Male dose-response requires further optimization to achieve more durable therapeutic effects.

The sex-specific therapeutic response to TNFR2 agonism parallels emerging evidence of sexually divergent TNF receptor signaling. We and others have demonstrated sex-dependent TNFR2 effects in preclinical models of neuropathic pain ^34^, multiple sclerosis ^40^ which investigated both male and female mice and showed that females consistently showed more robust responses than males. In these preclinical models, this sex-dependent response has been attributed to TNFR2-mediated expansion of regulatory T cells and downstream anti-inflammatory signaling. Based on established mechanisms, we hypothesize that similar TNFR2-driven Treg expansion may contribute to the modulation of fibrotic and immune-regulatory pathways in diastolic dysfunction, a possibility to explore in future studies. However, the predominance of contractile and ion-channel dysregulation in males may render immune modulation alone less effective. Direct comparison between TNFR2-agonism and current standard-of-care therapies remains necessary to position this approach within the existing therapeutic landscape. These findings underscore the importance of sex as a biological variable in both preclinical and clinical HF research and highlight TNFR2 agonism as a potential sex-informed therapeutic strategy.

## Conclusions

In conclusion, this study found that TNFR2 activation prevented and reversed diastolic dysfunction in female mice, whereas male mice showed an early benefit at higher doses that diminished over time. Mechanistically, TNFR2 stimulation modulated cardiac inflammation and pathological remodeling. These findings establish TNFR2 agonism as a potential sex-specific therapeutic strategy for inflammation-driven CVD in a novel model of OA-induced cardiovascular dysfunction.

## Supporting information

Supplementary Table 1: List of Gene for Figure 4

Graphical Abstract Publication Licence

## Acknowledgments

This work was supported by the National Institutes of Health (NIH Grants R01NS051709 to J.R.B.) and R01HL14729 to D.M. and M.W.K and the Department of Defense (DoD Grant W81XWH2110599 to J.R.B.). Moreover, this study was funded by resano GmbH to J.R.B. **Authors’ Contribution:** Conceptualization: PP and J.R.B; Writing-Original draft preparation: PP and KT; OA surgery and pain assessment: PP; Data arrangement: KT and PP. ECHO and WB: KT; Histology and analysis: KT, MD, LB, and LS. RNA-seq data analysis: AM and KT; Optical Mapping: RR, MD, and M.W.K. Graphical abstract: M.E (66) and P.P.; Writing, review, and editing: all authors have contributed.

\# Pranav Prasoon (PP) and Kelly Tammen (KT) are equal contributors for this manuscript.

## Conflict of interest

R.F. and M.E. are employees at resano GmbH.

J.R.B. received research funding from resano GmbH.

R.F., M.E., P.P. and J.R.B. are named as inventors on patent applications relating to selective TNFR2 agonists and medical uses thereof.

## Notes

### Summary of Updates

We revised Figures 2, 3, 4, and 6. We updated the scale bar for Histology and Echo. Also, we updated the Figure legend and references.

