## Supplementary Figure 1-3 for "TNFR2 Agonism as a Sex-Specific Therapy for Novel Osteoarthritis-Induced Cardiac Dysfunction"

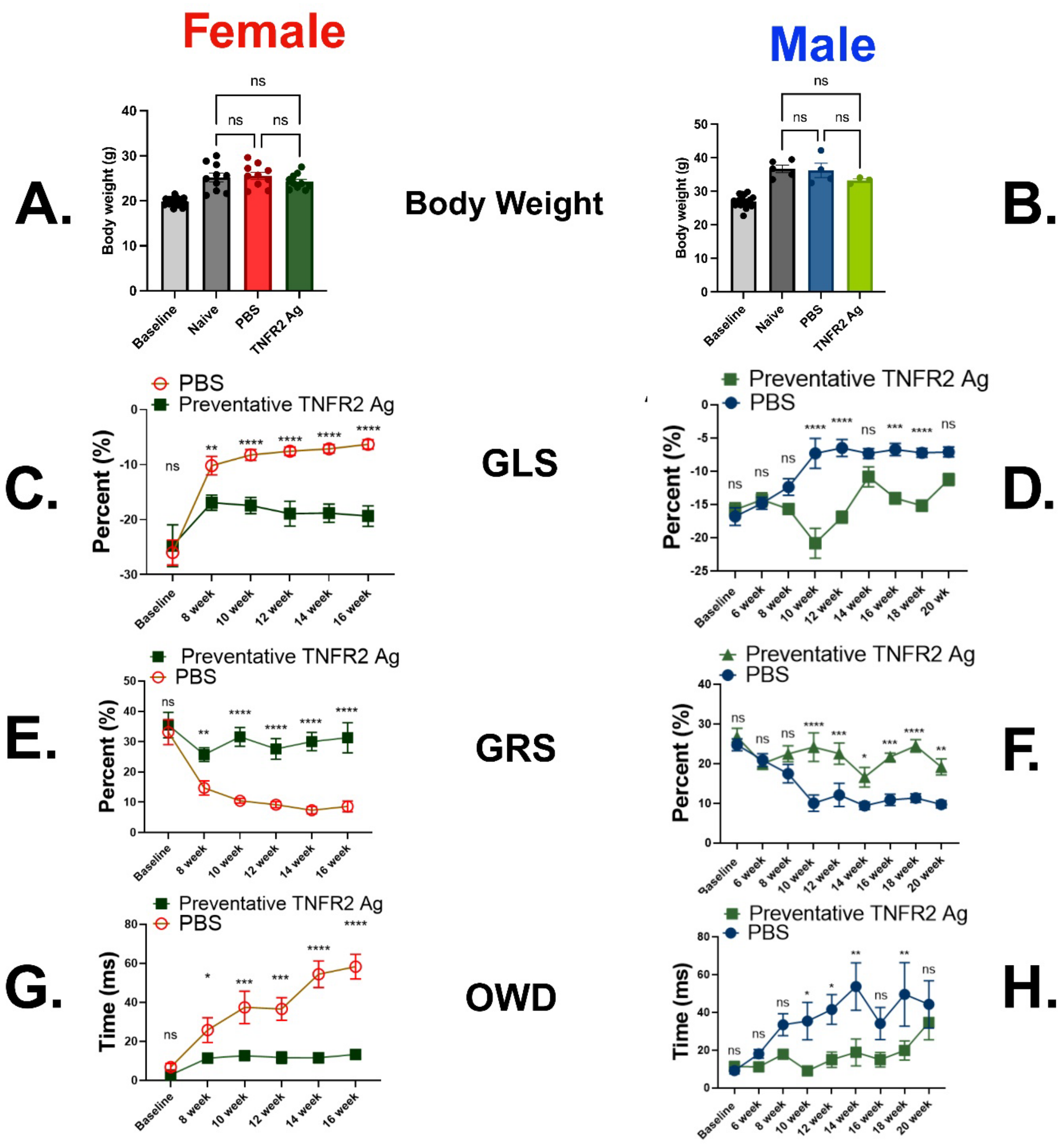


**Supplementary Figure 1: Preventive administration of TNFR2 agonist improved strain parameter in female (1 mg/kg) and male (10 mg/kg).**

**(A-B) Body Weight.** Body weight measurements at baseline and 16 weeks post-DMM for females and males show no differences between groups at 16 and 20 weeks, respectively, and increase in line with typical weight increases in Jackson C57BL6/J mice. **(C-D) Global Longitudinal Strain.** Global longitudinal strain (GLS) of male and female mice showed a significant increase in longitudinal strain in male and female vehicle-treated groups. Prophylactic TNFR2 Ag administration (1 mg/kg for females, 10 mg/kg for males) significantly improved strain in females at end timepoints and transiently in males. **(E-F) Global Radial Strain.** Global radial strain (GRS) of male and female mice showed a significant reduction of radial strain in male and female vehicle-treated groups. Prophylactic TNFR2 Ag administration significantly improved strain in female and male mice. **(G-H) Outer Wall Delay (OWD).** Outer wall delay in male and female mice showed a significant increase in vehicle-treated groups. Prophylactic TNFR2 Ag administration significantly improved the delay in females through end timepoints, while improving the delay transiently in males. Data are presented as mean ± S.E.M.; statistical analysis by one-way ANOVA with post hoc multiple comparisons. *$P < 0.05$, **$P < 0.01$, ***$P < 0.001$. Group sizes: n=8–10 for echocardiographic measures;

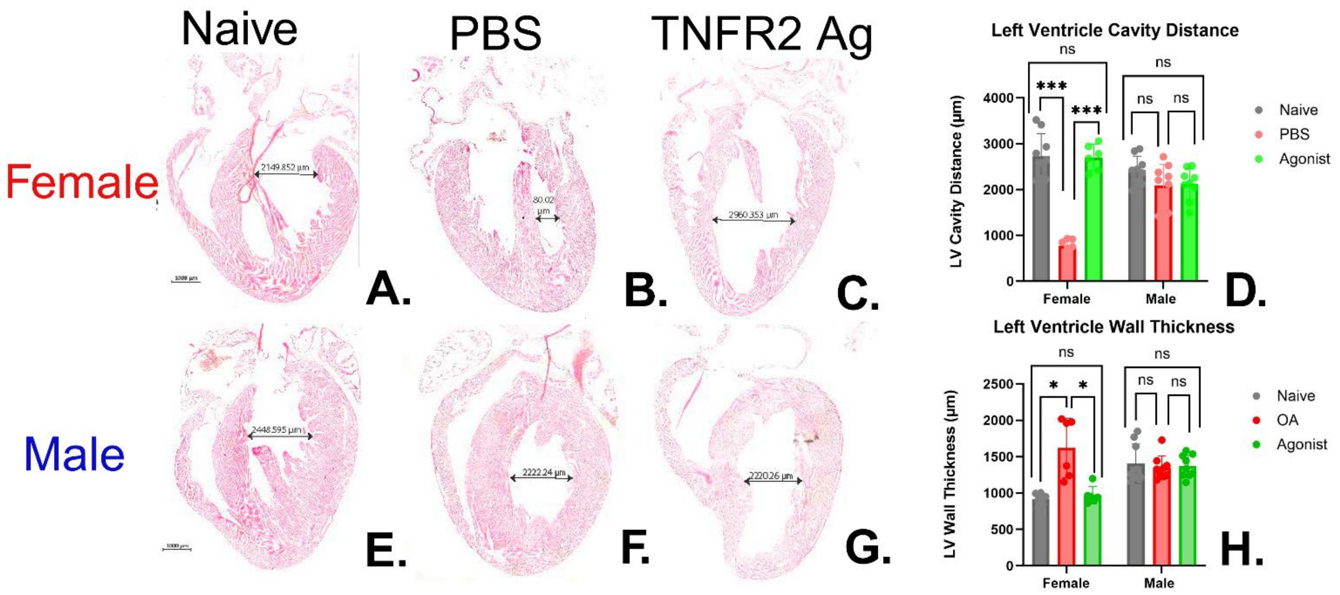


**Supplementary Figure 2: OA-induced distinct cardiac dysfunction in males and females. While TNFR2 agoinst (1mg/kg) was effective in females, not males.**

**(A-G)** Representative H&E Heart Stains. Representative hematoxylin and eosin (H&E) stained heart sections from naïve and OA mice demonstrate sex-specific patterns of cardiac hypertrophy. Naïve and OA female hearts, respectively. Naïve and OA male hearts, respectively. Female OA hearts exhibit concentric hypertrophy, characterized by reduced left ventricular (LV) chamber diameter and increased wall thickness, consistent with diastolic dysfunction. In contrast, male OA hearts exhibit eccentric hypertrophy; however, we did not observe changes in LV chamber diameter or ventricular wall thickness, despite the presence of an enlarged LV chamber diameter and thinner ventricular walls. This aligns with systolic dysfunction. However, TNFR2 agoinst (1 mg/kg) was effective in females, not males. **(D-H)** Quantification of LV chamber diameter and wall thickness from histological sections, displayed as bar diagrams, further highlights the sex-specific patterns of concentric versus eccentric hypertrophy. Scale bars = 1000 µm. Data are presented as mean ± S.E.M.; statistical analysis by one-way ANOVA with post hoc multiple comparisons. *P < 0.05, ***P < 0.001. Group sizes: n=6–8 for histology.

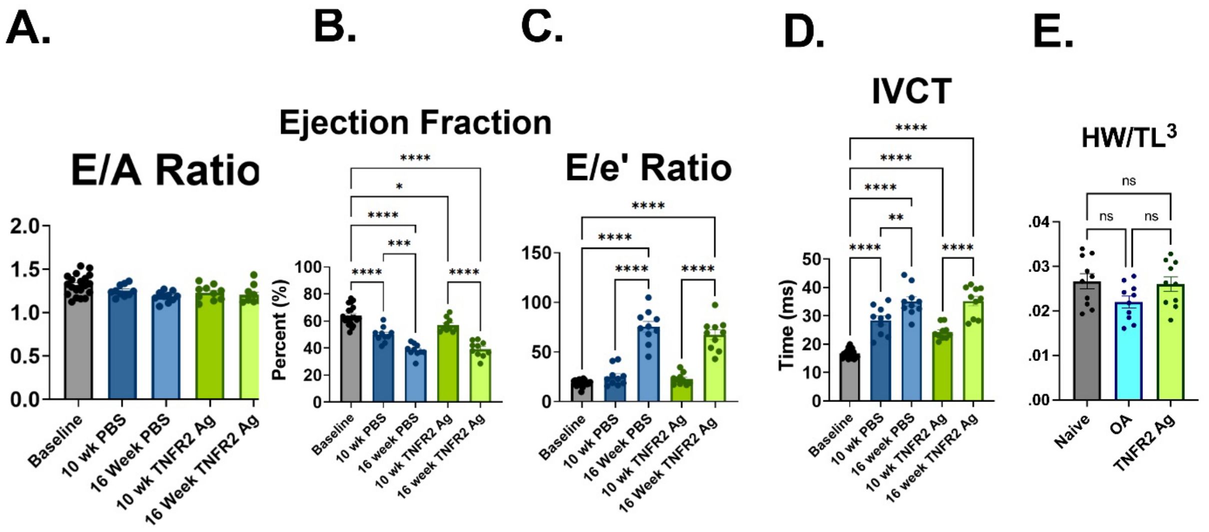


**Supplementary Figure 3: A low dose of TNFR2 agonist (1 mg/kg) does not effectively mitigate systolic dysfunction.**

(**A-D) Male cardiac function.** Quantitative analyses showed that males maintained the E/A ratio **(A)** at 10- and 16-weeks post-OA compared to baseline, but developed progressive systolic dysfunction characterized by increases in ejection fraction **(B)** and E/e′ ratios **(C)**. **(D)** Contraction abnormalities were evident with significantly prolonged isovolumetric contraction time (IVCT), and the TNFR2 Ag-treated mice at 1 mg/kg showed no difference from naïve controls at 10 or 16 weeks. **(E)** Male vehicle-treated mice exhibited no change in heart weight-to-tibia length ratio ($HW/TL^3$) when compared to naïve controls at 16 weeks. Data are presented as mean ± S.E.M.; statistical analysis by one-way ANOVA with post hoc multiple comparisons. *P < 0.05, ***P < 0.001. Group sizes: n=8–10 for histology.
